# Identification and capsular serotype sequetyping of *Streptococcus pneumoniae* strains

**DOI:** 10.1101/415422

**Authors:** Lucia Gonzales-Siles, Francisco Salvà-Serra, Anna Degerman, Rickard Nordén, Magnus Lindh, Susann Skovbjerg, Edward R. B. Moore

**Author notes:** Corresponding author E-mail address (LG) Post address: Guldhedsgatan 10A 41346 Gothenburg, Sweden.

## Abstract

Correct identification of *Streptococcus pneumoniae* (*pneumococcus*) and differentiation from the closely related species of the Mitis group of the genus *Streptococcus*, as well as serotype identification, is important for monitoring disease epidemiology and assessing the impacts of pneumococcal vaccines. In this study, we assessed the taxonomic identifications of 422 publicly available genome sequences of *S. pneumoniae, S. pseudopneumoniae* and *S. mitis*, using different methods. Identification of *S. pneumoniae*, by comparative analysis of the *groEL* partial sequence, was possible and accurate, whereas *S. pseudopneumoniae* and *S. mitis* could be misclassified as *S. pneumoniae*, suggesting that *groEL* is unreliable as a biomarker for differentiating *S. pneumoniae* from its closest related species. The genome sequences of *S. pneumoniae* and *S. pseudopneumoniae* fulfilled the suggested thresholds of average nucleotide identity (ANI), i.e., > 95% genome sequence similarity to the sequence of respective type strains for identification of species, whereas none of the *S. mitis* genome sequences fulfilled this criterion. However, ANI analyses of all sequences *versus* all sequences allowed discrimination of the different species by clustering, with respect to species type strains. The *in silico* DNA-DNA distance method was also inconclusive for identification of *S. mitis* genome sequences, whereas presence of the “Xisco” gene proved to be a reliable biomarker for *S. pneumoniae* identification. Furthermore, we present an improved sequetyping protocol including two newly-designed internal sequencing primers with two PCRs, as well as an improved workflow for differentiation of serogroup 6 types. The proposed sequetyping protocol generates a more specific product by generating the whole gene PCR-product for sequencing, which increases the resolution for identification of serotypes. Validations of both protocols were performed with publicly available *S. pneumoniae* genome sequences, reference strains at the Culture Collection University of Gothenburg (CCUG), as well as with clinical isolates. The results were compared with serotype identifications, using real-time Q-PCR analysis, as well as the Quellung reaction or antiserum panel gel-precipitation. Our protocols provide a reliable diagnostic tool for taxonomic identification as well as serotype identification of *S. pneumoniae*.

## INTRODUCTION

*Streptococcus pneumoniae* (pneumococcus) causes invasive and non-invasive disease, including pneumonia, meningitis, sepsis, otitis media, sinusitis, among others, particularly in children under the age of 5 years and the aged (Johnson et al., 2010), leading to approximately a million deaths annually in children aged less than 5 years, globally (Collaborators, 2017). A characteristic feature and the main virulence factor of *S. pneumoniae* is the polysaccharide capsule that enables the bacterium to evade host defence mechanisms (Nelson et al., 2007) and which is the basis for epidemiological categorization of pneumococcal isolates and strains into serotypes and serogroups (Geno et al., 2015). To date, 97 different capsular serotypes within 46 serogroups of *S. pneumoniae* have been identified on the basis of the biochemical structure of the capsular polysaccharide (Geno et al., 2015).

Several pneumococcal vaccines, which differ according to the polysaccharide capsule composition, have been developed. The first pneumococcal conjugate vaccine (PCV), licensed in 2000, covered 7 serotypes (PCV7: 14, 6B, 19F, 23F, 4, 9V, 18C) (Hicks et al., 2007), followed by PCV10 (PCV7 serotypes plus serotypes 1, 5, and 7F) in 2009 (Esposito and Principi, 2015), PCV13 (PCV10 serotypes plus serotypes 3, 6A, and 19A) in 2010 (Geno et al., 2015). A 15-valent conjugate vaccine is currently in clinical trials, and includes also serotypes 22F and 33F (LeBlanc et al., 2017). The pneumococcal polysaccharide vaccine (PPSV23) protects against 23 different capsular types (1, 2, 3, 4, 5, 6B, 7F, 8, 9N, 9V, 10A, 11A, 12F, 14, 15B, 17F, 18C, 19A, 19F, 20, 22F, 23F and 33F), and covers a high percentage of the types found in pneumococcal bloodstream infections. The vaccine is widely used for adults who are considered to be at high risk, as well as in children older than 2 years and at increased risk for pneumococcal disease (Diao et al., 2016). The use of the conjugate vaccines have significantly reduced the burden of pneumococcal disease in many populations. However, since vaccine introduction, “serotype replacement” has been observed, with increases in the proportions of invasive and non-invasive disease caused by pneumococcal serotypes not covered by the vaccines (Hicks et al., 2007; Weinberger et al., 2011).

*S. pneumoniae* serotype 6C is an example of an opportunistic increase in an infectious pneumococcus through serotype replacement. Serotype 6C was described as a newly-recognized serotype in 2007 (Mavroidi et al., 2004) and appears to have been rare in pre-vaccination populations. However, since the introduction of PCV7, the incidence of serotype 6C in disease and carriage has increased in diverse populations, worldwide (Loman et al., 2013). PCV7 contains polysaccharide from the 6B serotype capsule and PCV13 later included capsular polysaccharide of serotype 6A, although current vaccines do not extend protection to serotype 6C, which likely has promoted the observed serotype replacement (Park et al., 2008). Such serotype transitions demonstrate the importance of maintaining surveillance programs and clinical protocols that are able to respond to the evolutionary plasticity of infectious disease.

The classical serotyping method, the Quellung reaction, is based on the reaction of serotype-specific antisera with the corresponding capsule (Neufeld F, 1910). This method is time-consuming and costly, requiring live, cultivable bacteria, and a high degree of expertise, to the point that few laboratories are able to carry out the analyses. During the last decade, the nucleotide sequences of the capsule polysaccharide synthesis (CPS) loci (*cps*), harbouring the genes responsible for synthesis of the pneumococcal cell polysaccharide capsule, have been determined for all known serotypes. Accordingly, DNA amplification-based methods targeting specific capsular synthesis genes that allow differentiation of the serotypes have been developed, i.e., sequential multiplex PCR and sequential real-time Q-PCR (Pai et al., 2006; Varghese et al., 2017). Recently, a PCR-amplification and DNA-sequence-based typing method, ‘sequetyping’, was described targeting the regulatory gene, *cpsB*, with a single multiplex PCR, enabling the amplifications of 84 serotypes and sequencing of PCR-products, differentiating 46 of the 93 serotypes recognized at that time (Leung et al., 2012).

As an important practical step, before initiating pneumococcal serotype identification, it is critical to confirm the identification of *S. pneumoniae* and differentiate it from the other species of the Mitis group of the genus *Streptococcus* (Kawamura et al., 1995). The most closely-related species of *S. pneumoniae* are *S. pseudopneumoniae* and *S. mitis*. Sequencing of the 16S rRNA genes identifies a cytosine nucleotide at position 203 as a pneumococcal sequence signature, with an adenosine residue in all other species of the Mitis group (Scholz et al., 2012). Partial sequence determinations of individual metabolic ‘housekeeping’ genes, as a multi-locus sequence analysis (MLSA) (Bishop et al., 2009), continue to be widely used for identifying strains at the species level; for *Streptococcus*, *groEL*, *gyrB*, *rpoB* and *sodA* have been described as biomarker “housekeeping” genes for identification of the species in the *Streptococcus* genus (Glazunova et al., 2009; Hoshino et al., 2005; Kawamura et al., 1999; Teng et al., 2002). Additionally, the recently described “Xisco” gene, which has been demonstrated to be a unique biomarker for *S. pneumoniae,* provides a new approach for confirming specific differentiation between *S. pneumoniae* and its close relatives of the Mitis group (Salvà-Serra et al., 2017). Genome-based methods, such as average nucleotide identity (ANI) and *in silico* DNA-DNA hybridization, are gaining recognition as robust measurements of relatedness between strains, with potential in confirming phylogenetic and taxonomic relationships of bacterial identification (Konstantinidis and Tiedje, 2005; Meier-Kolthoff et al., 2013).

In this study, we present an improved workflow for pneumococcal serotype identification, including subtyping within serogroup 6 by sequetyping, as well as *S. pneumoniae* species confirmation and differentiation from closely related streptococcal species.

## METHODS

### Bacterial strains

One-hundred thirty-eight pneumococcus strains with identified serotypes were obtained from the Culture Collection University of Gothenburg, Gothenburg, Sweden (CCUG), where they were maintained in lyophilized state for long-term storage. The serotypes of these strains were determined at the Statens Serum Institut in Copenhagen, Denmark, by the Quellung reaction (Slotved et al., 2016), or at the Public Health Agency of Sweden, using an antiserum panel gel-precipitation protocol (Jauneikaite et al., 2015). Additionally, 50 strains, isolated from blood and cerebrospinal fluid samples during 2013 and 2014 and identified as *S. pneumoniae* at the Clinical Microbiological Laboratory, Sahlgrenska University Hospital, Gothenburg, Sweden, were included in the study. The strains isolated from clinical samples were stored in freeze-drying medium (Fry and Greaves, 1951), at −70 °C.

### Genome sequence data

A local database was created, including all genome sequences of *S. pneumoniae* (n=328) that were available in GenBank (Benson et al., 2017) on the 14th March 2015, plus the type strain *S. pneumoniae* NCTC 7465^T^ (GenBank accession number: LN831051) and all genome sequences that were available in GenBank on the 18 ^th^ May 2016 for 14 other species of the Mitis group (n=248): *S. pseudopneumoniae* (n=40), *S. mitis* (n=53), *S. australis* (n=2), *S. cristatus* (n=16), *S.dentisani* (n=2), *S. gordonii* (n=22), *S. infantis* (n=7), *S. massiliensis* (n=2), *S. oralis* (n=34), *S. parasanguinis* (n=29), *S. peroris* (n=1), *S. sanguinis* (n=33), *S. sinensis* (n=1) and *S. tigurinus* (n=6) (Jensen et al., 2016).

### DNA extraction

The stored strains from CCUG or clinical samples were inoculated onto Blood Agar plates with horse blood 5% (prepared at the Substrate Department, Clinical Microbiological Laboratory, Sahlgrenska University Hospital), and incubated overnight at 36 °C with 5% CO2. DNA was extracted, using a ‘heat-shock’ protocol (Welinder-Olsson et al., 2000). Briefly, an inoculating loop-full of bacterial biomass was suspended and incubated in 100 μL Tris-EDTA buffer and 15 μL lysostaphin 0.05 μM (Sigma-Aldrich, St. Louis, MO, USA) at 37 °C for 1 hour. Subsequently, 10 μL of Proteinase K (Sigma-Aldrich, St. Louis, MO, USA) were added and the suspensions were incubated for 2 hours at 56 °C. Finally, the samples were incubated at 95 °C for 10 minutes. After incubation, samples were centrifuged at 17,900 x *g*, for 10 min. The supernatant containing genomic DNA, was transferred to a new tube and stored at −20 °C.

For multiplex PCR analyses, bacterial DNA was extracted, using a MagNA Pure LC instrument (Roche Diagnostics, Mannheim, Germany) and a Total Nucleic Acid Isolation kit (Roche Diagnostics, Mannheim, Germany). The extracted DNA was eluted in 100 μl of elution buffer, and stored at −20 °C, until real-time multiplex PCR-assays were performed.

### Taxonomic identifications

Identifications of reference strains and strains isolated from clinical samples were determined by PCR-amplification and sequence analysis of partial (757 bp) *groEL* gene, using primers, StreptogroELd and StreptogroELr, as previously described (Glazunova et al., 2009). PCR-products were purified and sequenced (GATC Biotech AG, Constance, Germany). The sequences were compared with the *groEL* partial sequences of the type strains of the 20 validly published species of the Mitis group of the genus *Streptococcus*, using BioNumerics software platform, version 7.5 (Applied Maths, Sint-Martens-Latem, Belgium). A strain was assigned to a given species if the partial *groEL* sequence similarity value was above 96%. The strains were also analysed for presence of the “Xisco” gene, using amplification-primers, Spne-CW-F2 and Spne-CW-R, according to Salvà-Serra *et al*. (2017).

The taxonomic status of the 422 genome sequences of *S. pneumoniae*, *S. pseudopneumoniae* and *S. mitis* included in the local data base were assessed by determining average nucleotide identity, based on BLAST (ANIb) (Goris et al., 2007), using JSpeciesWS (Richter et al., 2016), against the reference genome sequences of the type strains of the different species. Additionally, the matrix obtained from ANIb similarities of all vs. all genome sequences was used to construct an ANIb-based dendrogram, according to Gomila *et al*. (2015). Briefly, the matrix of ANIb values was used, applying Pearson’s distance correlation and an average linkage construction (UPGMA hierarchical clustering), using PermutMatrix software (Caraux and Pinloche, 2005). Finally, *in silico* DNA-DNA distance values were calculated, using the Genome-to-Genome Distance Calculator (GGDC), (ggdc.dsmz.de) (Meier-Kolthoff, 2013) and the recommended BLAST+ method. The GGDC results shown are based on the recommended formula 2 (sum of all identities found in high-scoring segment pairs (HSPs), divided by the overall HSP length), which is independent of genome size and is, thus, robust when using draft genomes.

### Modified Sequetyping protocol

The sequetyping protocol was based on analysis of the capsule polysaccharide synthesis *cpsB* region (Leung et al., 2012). In order to obtain sufficient quality for the entire 1,061 bp segment, two internal primers were designed, wzh-mid-F and wzh-mid1-R, generating two partly overlapping sequences (Figure 1). The reaction mixture for PCR-assays comprised 0.1 to 10 ng of DNA template, 1X Taq PCRMasterMix (Qiagen, Hilden, Germany), 1 μM concentration of each amplification-primer, in a total volume of 25 μL. Primer sequences are listed in Supplementary Table 1. PCR-amplification was achieved, with an initial cycle of 5 min denaturation at 95°C and 30 cycles of 30 s at 95°C for denaturation, 30 s at 55°C for primer-annealing and 90 s at 72°C for primer-extension, with a final extension step at 72°C for 5 min. Amplicons were analysed by electrophoresis in 1% agarose gel. Sequencing reactions were performed using the four primers (Figure 1).

**Figure 1.**
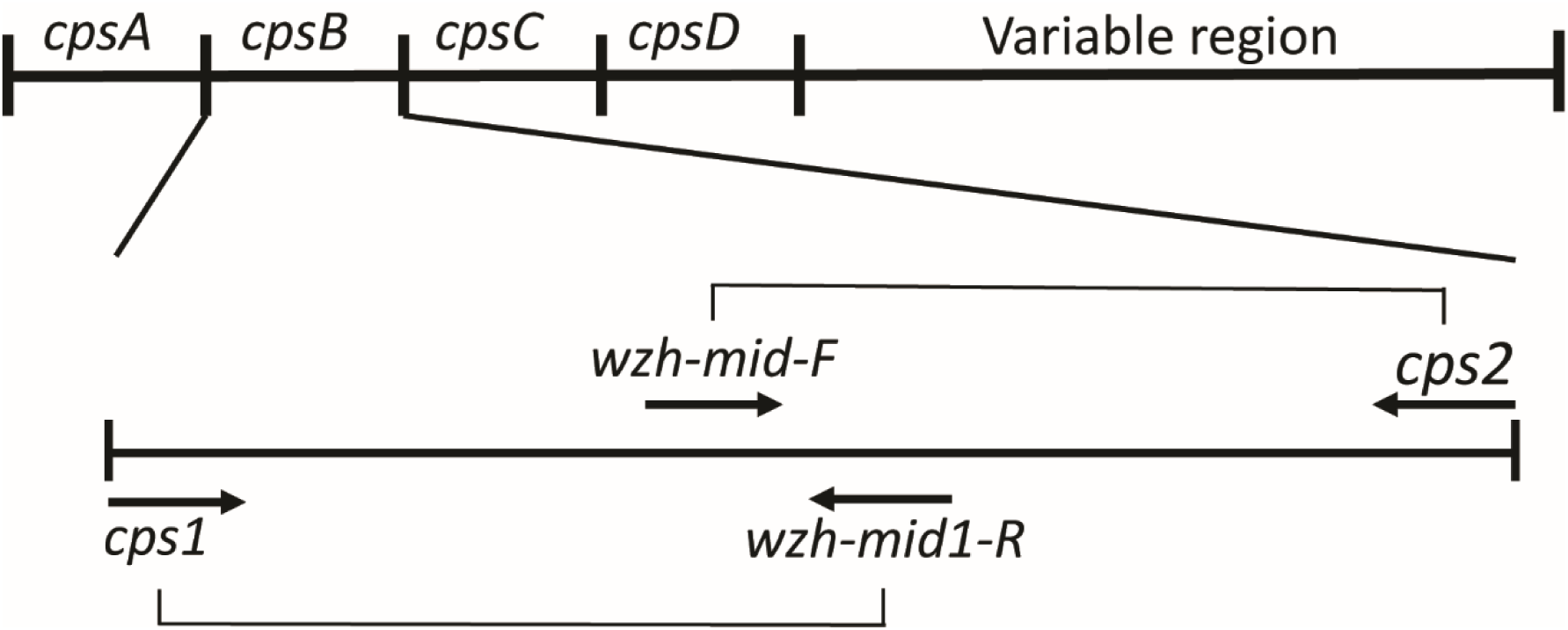
Schematic representation of the targeted *cpsB* gene in the conserved region with *cpsA*, *cpsB*, *cpsC* and *cpsD* within the CPS loci of *Streptococcus pneumoniae* for amplification, using two primer pairs.

A database of reference sequences was created for comparative analyses, including the sequences for each serotype listed by Bentley et al., (2006), as well as the complete *cpsB* region sequences extracted from the 329 *S. pneumoniae* genome sequences. The PCR-amplicon nucleotide sequences were analysed by similarity analysis, using BioNumerics software platform, version 7.5 (Applied Maths, Sint-Martens-Latem, Belgium). A strain was assigned to a given serotype if the similarity value was higher than 99% and the similarity of the second highest match was, at least, 1% lower. If the similarity value was shared between two or more serotypes, it was reported as multiple-matched serotypes.

BLASTN analyses of the *cpsB* region were also performed, with respect to the 248 non-pneumococcal genome sequences, in order to determine if this region is present in other species of the Mitis group. Only BLAST hits with an E-Value lower then 10e-5 were considered significant. Additionally, the four sequetyping primers were also analysed, using BLASTN against the 248 non-pneumoniae genome sequences. Only matches covering the entire primer length (100% coverage) and maximum 2 mismatches were considered to be positive.

### Serotype identification by multiplex real-time PCR

A multiplex real-time PCR, able to detect 40 different serotypes, was developed and applied. The assay is based on a protocol published by Centers for Disease Control and Prevention (CDC) (da Gloria Carvalho et al., 2010), and is similar to the real-time PCR subsequently developed by CDC (Pimenta et al., 2013). A complete list of primers and probe sequences can be found in Supplementary Table 2. The multiplex real-time PCR was performed in 384-well format in a Quant Studio 6 Flex (Applied Biosystems, Carlsbad, CA). Each PCR consisted of a 20 μl reaction volume, including 4 μl of template DNA, along with 1 μM of each of the forward and reverse primers, 0.85 μM of the probe, 10 μl of 2X Universal Master Mix for DNA targets (Applied Biosystems, Foster City, CA, USA) and RNAase-free water. The Tecan Freedom EVO PCR setup workstation (Life Sciences, Männedorf, Switzerland) was used to prepare the PCR assays in a 384-well plate. The reaction conditions were as follows: one initial cycle at 46°C for two minutes, followed by denaturation at 95°C for 10 minutes and 45 amplification cycles of 95°C for 15 s and 58°C for one minute. Each multiplex performance was evaluated, using an internal control (*cpsA*) to verify the presence of pneumococcal DNA in the sample, as well as two pUC57 plasmids containing each PCR target amplicon for all serotype systems.

## RESULTS

### Classification of database sequences

In order to re-evaluate the classifications of the genome sequences of *S. pneumoniae*, *S. pseudopneumoniae* and *S. mitis* in the genome sequence database, ANIb analyses were performed, wherein each genome sequence was compared to that of the type strain of each species of the Mitis group and by comparisons of all genomes to all. The analyses showed that all 328 strains (type strain excluded) listed as *S. pneumoniae* in the GenBank database were correctly identified as *S. pneumoniae*, i.e., had similarity values greater than 95% (Rosselló-Móra and Amann, 2015). Of them, 24 sequences exhibited ANIb similarity values ≥ 99% to the sequence of the type strain, 271 strains exhibited ≥ 98% similarity and 33 strains exhibited ≥ 97% similarity. By comparison, only 9 of the 39 sequences from strains listed as *S. pseudopneumoniae* (type strain excluded) in GenBank exhibited ANIb values ≥ 95%, whereas 48 of the 52 sequences from strains listed as *S. mitis* (type strain excluded) exhibited ANIb values below 95% and only four strains had ANIb values between 95-96%, indicating a significant number of misclassifications of strains for which genome sequence data had been submitted to GenBank.

Additionally, cluster analyses was done, using the calculated ANIb similarity values for all strains against all, for *S. pseudopneumoniae* and *S. mitis*, including, as well, the type strains of the other 13 species of the Mitis group included in the genomes database. With this analysis, a dendrogram was generated, to visualize the relationships among the strains, with respect to the type strains of the different species (Supplementary Figure 1). In the cases of *S*. *pneumoniae*, all genome sequences clustered most closely with the type strain of *S. pneumoniae*, confirming the taxonomic designations for the genome sequences. However, only nine of the 39 genome sequences listed as *S. pseudopneumoniae* and thirty-six of the 52 genome sequences listed as *S. mitis* in the database clustered in proximity to the type strain of the respective species and were, therefore, taxonomically designated as *S. pseudopneumoniae* and *S. mitis*, while the remaining 46 strains clustered closer to other species.

The GGDC analyses, comprising comparison with type strains for each species, showed that all *S. pneumoniae* genome sequences had *in silico* hybridization values higher than 70%, confirming their taxonomic identities. The GGDC analyses for *S. pseudopneumoniae* matched the results obtained by ANIb analysis (Table 1), whereas only one of the genome sequences of *S. mitis* exhibited a hybridization value higher than 70%; for the rest of the genome sequences, the *in silico* DNA-DNA hybridization values were lower than 70% and inconclusive for confirming species-level identifications (Table 2).

**Table 1.** Taxonomic identification of the correctly-classified *Streptococcus pseudopneumoniae* strains from GenBank.

| GenBank<br>strain | ANiB all Vs all | groEL similarity |  |  |  |  |  | ANiB similarity |  |  | GGDC<br>(%) |
| --- | --- | --- | --- | --- | --- | --- | --- | --- | --- | --- | --- |
|  |  | 1st match | % | 2nd match | % | 3th match | % | *S. pseudopneumoniae* | *S. pneumoniae* | *S. mitis* |  |
| 1321 | *S. pseudopneumoniae* | *S. pseudopneumoniae* | 100 | *S. mitis* | 95.8 | *S. pneumoniae* | 94.1 | 98.4 | 94.2 | 91.8 | 86.6 |
| 22725 | *S. pseudopneumoniae* | *S. pneumoniae* | 99.6 | *S. pseudopneumoniae* | 93.9 | *S. mitis* | 93.4 | 96.9 | 94.1 | 91.9 | 74.9 |
| 276-03 | *S. pseudopneumoniae* | *S. pneumoniae* | 99.5 | *S. pseudopneumoniae* | 93.8 | *S. mitis* | 93.3 | 97 | 94.1 | 91.9 | 75.6 |
| 338-14 | *S. pseudopneumoniae* | *S. pneumoniae* | 99.6 | *S. pseudopneumoniae* | 93.9 | *S. mitis* | 93.4 | 97 | 94.1 | 91.8 | 75.5 |
| 5247 | *S. pseudopneumoniae* | *S. pseudopneumoniae* | 96.4 | *S. mitis* | 95.7 | *S. pneumoniae* | 95 | 96.5 | 94.1 | 91.9 | 72.3 |
| 61-14 | *S. pseudopneumoniae* | *S. pneumoniae* | 99.6 | *S. pseudopneumoniae* | 93.9 | *S. mitis* | 93.4 | 96.8 | 94 | 91.6 | 75.5 |
| G42 | *S. pseudopneumoniae* | *S. pneumoniae* | 99.1 | *S. pseudopneumoniae* | 94.5 | *S. mitis* | 93.9 | 96.9 | 94 | 91.8 | 75.3 |
| IS7493 | *S. pseudopneumoniae* | *S. pneumoniae* | 99.6 | *S. pseudopneumoniae* | 93.9 | *S. mitis* | 93.4 | 96.8 | 94 | 91.9 | 73.8 |
| SK674 | *S. pseudopneumoniae* | *S. pseudopneumoniae* | 100 | *S. mitis* | 95.8 | *S. pneumoniae* | 94.1 | 98.8 | 94.1 | 91.9 | 89.9 |

The results show the 9 of the 39 genome sequences classified as *S. pseudopneumoniae* at the GenBank database that were confirmed to their taxonomical identity as *S. pseudopneumoniae* by ANiB and GGDC analyses.

**Table 2.**
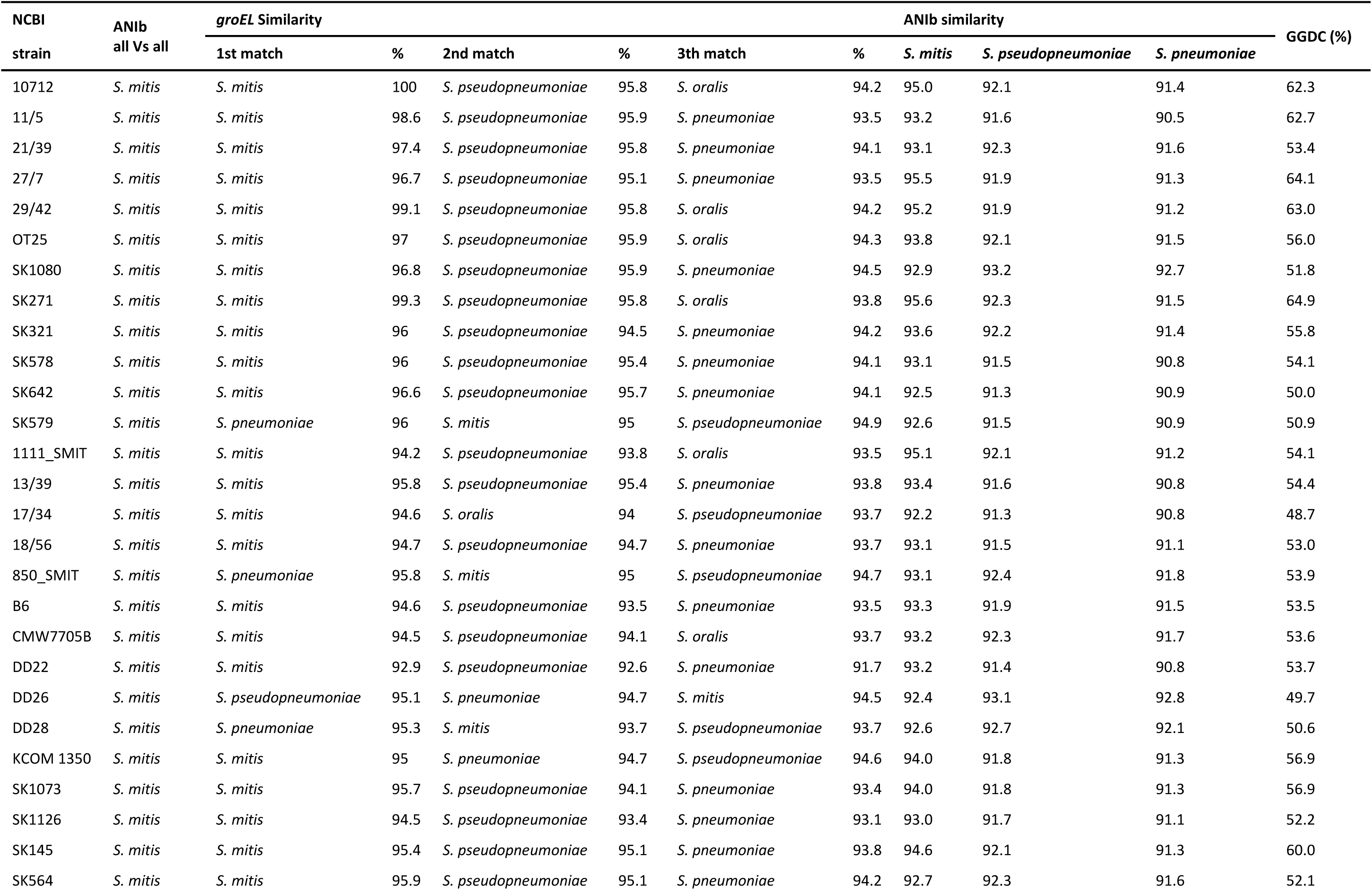
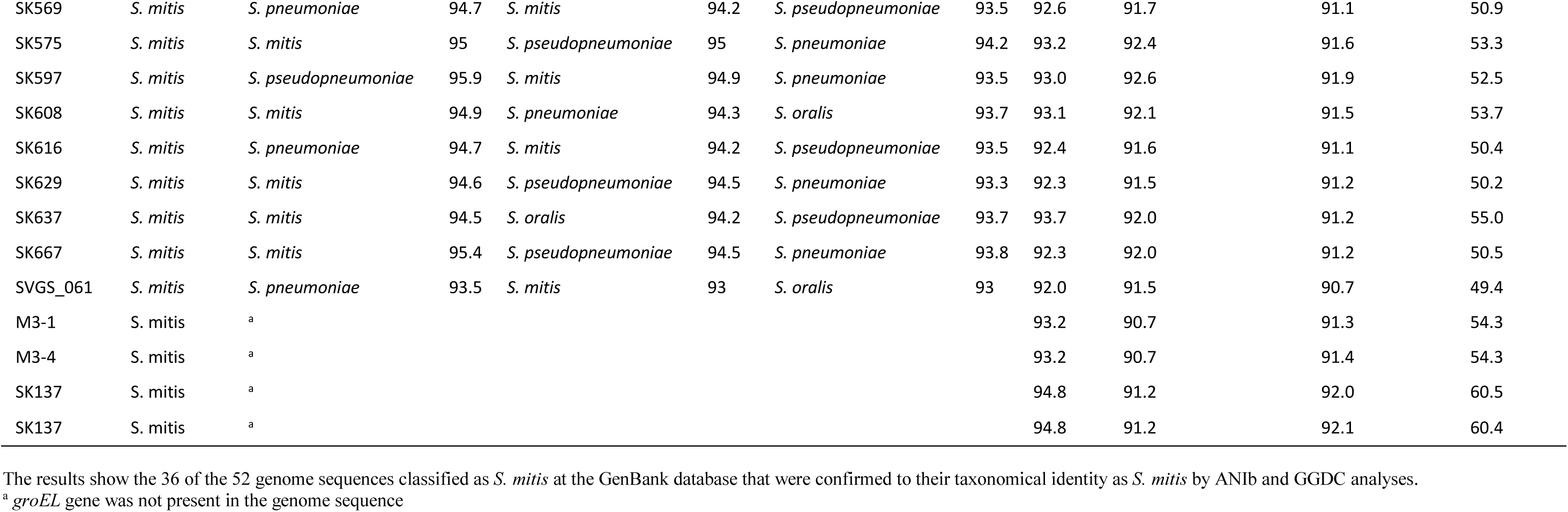
Taxonomic identifications of the correctly-classified *Streptococcus mitis* strains from GenBank.

Partial *groEL* sequence analyses using the region of the gene suggested by Glazunova *et al*., (2009) were also performed. The 757 bp *groEL* sequence was extracted from all genome sequences of *S. pneumoniae, S. mitis* and *S. pseudopneumoniae* and similarity values of the sequences were calculated, with respect to the type strains of the 14 species of the Mitis group of *Streptococcus*. By this analysis of partial *groEL* sequences, with sequence similarities above 96% (cut-off value) with the type strains of the respective species, all 328 genome sequences listed as representing *S. pneumoniae* genomes (type strain excluded) were identified as *S. pneumoniae*, whereas only three of 39 genomes listed as *S. pseudopneumoniae* (type strain excluded) were identified as *S. pseudopneumoniae* (Table 1), and 28 of the 52 sequences listed as *S. mitis* (type strain excluded) were identified as *S. mitis*. In four of the strains, the *groEL* gene could not be found in the genome sequence (Table 2). The classifications of the remaining 17 genome sequences were ambiguous, with non-definitive similarity values for *S. mitis*, as well as *S. pneumoniae* and *S. pseudopneumoniae*, allowing no clear species-level identifications (Table 2). Finally, the “Xisco” gene was not detected in any genome sequence listed as *S. mitis* or *S. pseudopneumoniae*, whereas the “Xisco” gene was present in all genome sequences identified as *S. pneumoniae*. A summary of the results for the genome sequences that were taxonomically incorrect but listed in GenBank as *S. pseudopneumoniae* and *S. mitis* are presented in Supplementary Table 3.

Based on these results, discrepancies were observed when comparing the results of identifications of genome sequences obtained by genome sequence ANIb analysis and results obtained by partial *groEL* sequencing, suggesting that *groEL* may not be as reliable a marker as anticipated for identification of the closely related species of the Mitis group of the *Streptococcus*.

### Identification of *Streptococcus pneumoniae* in culture collections and clinical samples

In cultivated and isolated clinical strains (n=50) as well as in the 138 strains from the CCUG previously identified as *S. pneumoniae*, the “Xisco” gene was present in 100% of strains. Furthermore, *groEL* similarity values were observed to be greater than 98% in all strains and greater than 99% in two-thirds of the analysed strains, confirming by two independent techniques that the strains were correctly identified as *S. pneumoniae*.

### Serotype identification

The sequetyping technique of Leung *et al*. (2012) was modified by using two internal primers to generate two partially overlapping amplicons, representing the whole 1,017 bp *cpsB*-region (Figure 1). To assess its accuracy, sequetyping was evaluated *in silico* by analysing *cpsB* sequences that were extracted from the 329 genome sequences of *S. pneumoniae* in the local genome sequence database. The serotypes of 261 (80%) of these genomes were identified, with similarity values greater than 99% to a reference sequence. In 15 strains similarity matches were low and therefore they could not be assigned to the serotype, 8 strains exhibited best matches to serotypes 10B-10C (less than 97%) and 7 strains exhibited best matches to serotype 24F (less than 98%). The serotypes of 52 genomes could not be determined. Thirty-six of these genomes were previously described as ‘non-typeable’ pneumococci (Hathaway et al., 2004) and, therefore, the *cpsB* sequence was not present (Table 3). For the remaining 16 genomes, the serotypes could not be determined, due to low similarity values, with respect to the reference sequences.

**Table 3.** Serotypes identified by sequetyping, using *in silico* analysis of the 329 genome sequences of *S. pneumoniae* from GenBank included in our local database.

| Serotype | #strains |
| --- | --- |
| 1 | 33 |
| 2/41A | 1 |
| 3 | 14 |
| 4 | 11 |
| 5 | 1 |
| 6 | 32 |
| 7A/7F | 7 |
| 8 | 2 |
| 9A/9V | 11 |
| 9N/9L | 2 |
| 10A | 3 |
| 10C | 1 |
| 10C/10F<sup>a</sup> | 8 |
| 11a/11D/18F | 6 |
| 12a/46 | 3 |
| 13/20 | 2 |
| 14 | 17 |
| 21 | 1 |
| 15A/15F | 4 |
| 16F | 1 |
| 17F/33C | 4 |
| 18B/18C | 3 |
| 19A | 48 |
| 19B | 1 |
| 19C | 2 |
| 19F | 28 |
| 22A/22F | 5 |
| 23A | 2 |
| 23F | 16 |
| 24F<sup>a</sup> | 7 |
| 33A/35A/33F | 2 |
| 35F/47F | 1 |
| NT | 36 |
| NI | 14 |

<sup>a</sup> match with low similarity value NT, non-typable NI, non-identified

BLASTN analyses of the *cpsB* region, with respect to the 248 non-pneumoniae genome sequences, gave 22 positive hits with E-Values lower than 10e-5 but with similarities ranging between 93 and 80% (Supplementary Table 4), suggesting that the *cpsB* region could also be present in other species of the Mitis group. Analysis of the probability for the four sequetyping primers to amplify among the 248 non-pneumoniae genome sequences considering a maximum of 2 mismatches showed that the PCR including the primers cps1-*wzh*-mid-R could lead to 2 positive amplifications, whereas the PCR reaction including the primers *wzh*-mid-F – cps2 could lead to 13 positive amplifications. However, the whole *cpsB* region will be expected to be amplified in only two cases, in *S. mitis* SK579 and *S. mitis* SK616 (Supplementary Table 4).

The sequetyping was applied on the 138 *S. pneumoniae* strains from the CCUG, for which the serotypes had previously been determined by the Quellung reaction or the antiserum panel gel-precipitation protocol. The determined sequences were analysed by BLAST searches, and similarities were recorded. A sequence was assigned to a specific serotype if the similarity value was greater than 99% and the next-best similarity match was, at least, 1% lower. In 140 strains (97%), the serotype by sequetyping matched the results obtained by the reference methods.

Discrepancies were observed for five strains: CCUG 1749 (17A) was identified as 10A; CCUG 5906 (36) was identified as 15B; CCUG 20653 (48) was identified as 6B; CCUG 27692 (15A) was identified as 19B; and CCUG 55117 (16A) was identified as 48.

Finally, the serotypes of 50 strains isolated from clinical samples were determined, using sequetyping, multiplex real-time PCR and antiserum panel gel-precipitation protocol performed at the Public Health Agency of Sweden. A serotype was identified by real-time PCR for 36 of the 50 strains (73%), whereas the serotypes were identified only by sequetyping for the remaining 14 strains (27%), showing serotypes that were not targeted by the real-time PCR assay. In all cases, the obtained results agreed with those obtained by the antiserum panel protocol (Table 4).

**Table 4.**
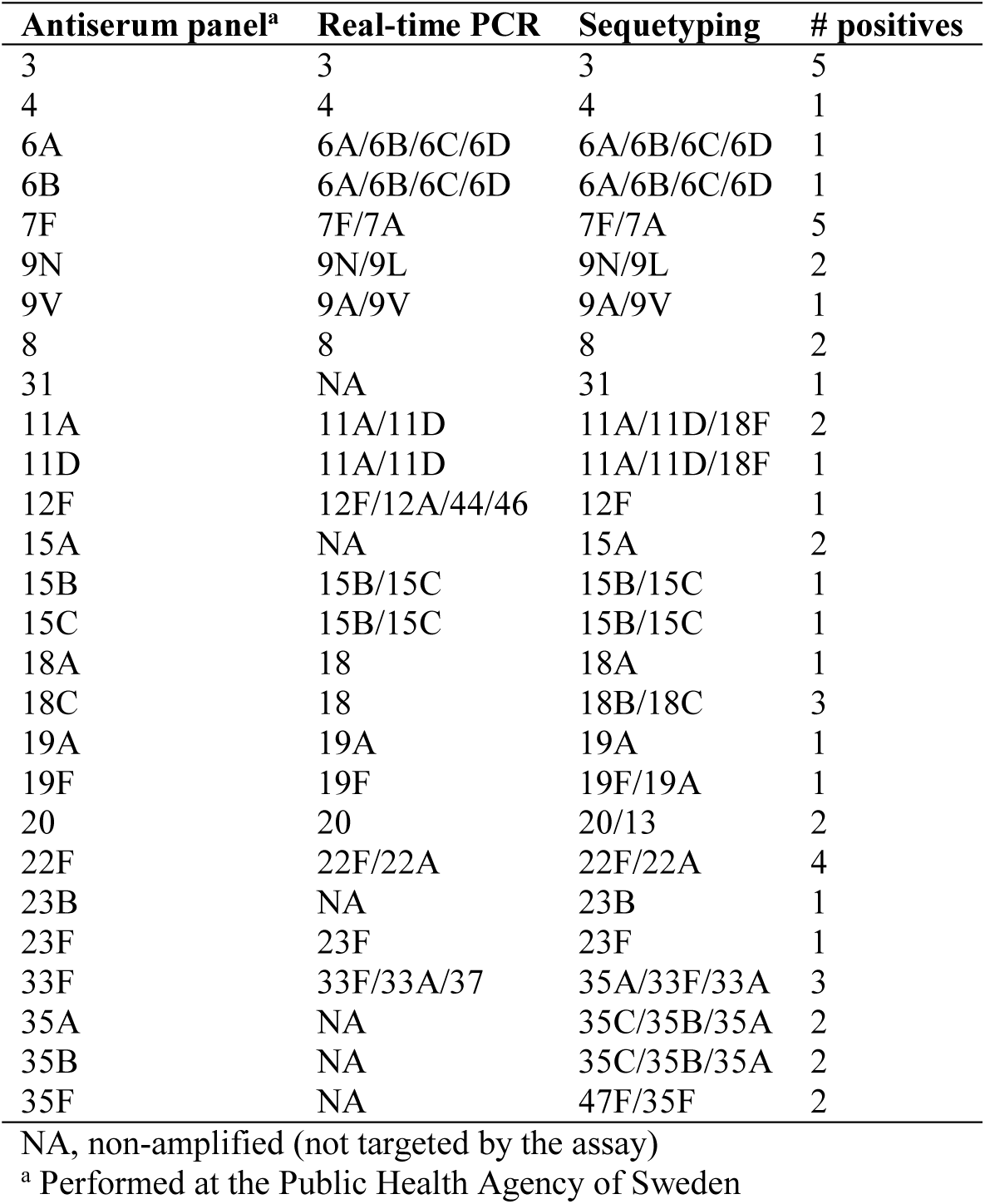
Serotype identification of strains isolated from clinical samples.

### Serogroup 6 differentiation

A dendrogram, based on the sequence of the entire *cpsB* region (1,017 bp) sequence from all strains classified as serogroup 6, was created (Supplementary Figure 2). The sequences did not form distinct clusters, indicating that serotype differentiation among serogroup 6 is not possible by sequetyping of this region and that an alternative method is needed. A DNA sequencing-dependent approach was used for differentiating the serotypes 6A, 6B, 6C and 6D; a schema of the suggested protocol is shown in Figure 2. Firstly, a PCR-amplification, using the primers, wciP374F (this study) and wciP-R (Jin et al., 2009), was performed and the PCR-product was sequenced, using primer, wciP374F. Primer sequences are listed in Supplementary Table 1. The sequence product allowed visualization of the single nucleotide polymorphism (SNP) that distinguishes serotype 6A/6C (guanine in position 584) and serotype 6B/6D (adenine in position 584). Subsequently, for differentiating serotype 6A and 6C, a second PCR, using primers, Del6Cwzy_Fv2 (this study) and Del6Cwzy_R (Jin et al., 2009), followed by Sanger sequencing, using primer Del6Cwzy_Fv2, was performed. A 6 bp deletion in the gene *wzy,* characteristic for serotype 6C, was detected. The *in silico* PCR analysis showed that it is possible to obtain PCR-products for serotypes 6A, 6B and 6C but not for serotype 6D (Table 5), although, the deletion was present only in serotype 6C. For differentiating serotype 6B from 6D, a PCR, using primers, wciN_6AB_F and wciN_6AB_R (this study), was performed. This PCR is shown to be unique for serotype 6B; thus, if the PCR-product was produced, the strain was assigned to serotype 6B, whereas, if the PCR was negative, the strain was assigned to serotype 6D. To finally confirm serotype 6D, two additional PCR-assays, targeting the *wciN*_*beta*_ region were performed, the first PCR, using primers, *wciN*_*beta*_S1/ *wciN*_*beta*_A2 (Jin et al., 2009), and the second, using primers, *wciN*_*beta*_S2/ *wciN*_*beta*_A1 (Jin et al., 2009). If at least one of the PCR-assays was positive, the strain was confirmed as serotype 6D. Details of the analysis are presented in Table 5.

**Table 5.** *In silico* identification of serogroup 6.

| *S. pneumoniae* | Accession | PCR 1<sup>a</sup> |  | PCR 2<sup>b</sup> |  | PCR<br>3<sup>c</sup> | PCR<br>4<sup>d</sup> | PCR<br>5<sup>e</sup> | Serotype |
| --- | --- | --- | --- | --- | --- | --- | --- | --- | --- |
| Strain | number | A/G | Serogroup | PCR | Deletion |  | S1-A2 | S2-A1 |  |
| 07AR0125 | AFBY01000000.1 | G | AC | + | + | - | + | + | 6C |
| 801 | AQTO01000000.1 | G | AC | + | - | + | - | - | 6A |
| 845 | AQTP01000000.1 | G | AC | + | - | + | - | - | 6A |
| 1488 | AQTQ01000000.1 | A | BD | + | - | - | - | - | 6B |
| 670-6B | CP002176.1 | A | BD | + | - | + | - | - | 6B |
| BHN191 | ASHN01000000.1 | A | BD | + | - | - | - | - | 6B |
| BHN237 | ASHO01000000.1 | A | BD | + | - | + | - | - | 6B |
| BHN418 | ASHP01000000.1 | A | BD | + | - | + | - | - | 6B |
| BHN427 | ASHQ01000000.1 | A | BD | + | - | + | - | - | 6B |
| BR1064 | AFBZ01000000.1 | G | AC | + | + | + | + | + | 6C |
| CCUG1350 | LQQG01000000.1 | A | BD | + | - | + | - | - | 6B |
| CDC1873-00 | ABFS01000000.1 | G | AC | + | - | + | - | - | 6A |
| EU-NP04 | AIKH01000000.1 | G | AC | + | + | + | + | + | 6C |
| GA02270 | AIKJ01000000.1 | G | AC | + | - | - | - | - | 6A |
| GA02506 | AILJ01000000.1 | A | BD | + | - | - | - | - | 6B |
| GA02714 | AIKK01000000.1 | G | AC | + | - | - | - | - | 6A |
| GA14373 | AILN01000000.1 | G | AC | + | - | - | - | - | 6A |
| GA17328 | AGPH01000000.1 | G | AC | + | - | - | - | - | 6A |
| GA17971 | AGPJ01000000.1 | G | AC | + | - | + | - | - | 6A |
| GA19077 | AGPK01000000.1 | G | AC | + | - | + | - | - | 6A |
| GA41437 | AGPN01000000.1 | G | AC | + | - | + | - | - | 6A |
| GA47033 | AGOA01000000.1 | G | AC | + | + | + | + | + | 6C |
| GA52306 | AGPZ01000000.1 | G | AC | + | + | + | + | + | 6C |
| GA60080 | ALCR01000000.1 | G | AC | + | + | + | + | + | 6C |
| GA60132 | ALCV01000000.1 | G | AC | + | + | + | + | + | 6C |
| GA60190 | ALCL01000000.1 | G | AC | + | + | + | + | + | 6C |
| NorthCarolina6A-23 | AGQL01000000.1 | G | AC | + | - | + | - | - | 6A |
| NP127 | AGQC01000000.1 | G | AC | + | - | + | - | - | 6A |
| SP6-BS73 | ABAA01000000.1 | G | AC | + | - | + | - | - | 6A |
| SPAR55 | ALCF01000000.1 | G | AC | + | - | + | - | - | 6A |
| WL400 | AVFA01000000.1 | G | AC | + | - | + | - | - | 6A |
| K15-99 | HQ662206.1 | A | BD | - | --- | - | + | + | 6D |
| K15-60 | HQ662205.1 | A | BD | - | --- | - | + | + | 6D |
| K15-17 | HQ662214.1 | A | BD | - | --- | - | + | + | 6D |
| K15-129 | HQ662208.1 | A | BD | - | --- | - | + | + | 6D |
| K15-115 | HQ662207.1 | A | BD | - | --- | - | + | + | 6D |
| K13-22 | HQ662215.1 | A | BD | - | --- | - | + | + | 6D |
| K13-110 | HQ662218.1 | A | BD | - | --- | - | + | + | 6D |
| K13-109 | HQ662217.1 | A | BD | - | --- | - | + | + | **6D** |
| K13-108 | HQ662216.1 | A | BD | - | --- | - | + | + | **6D** |
| B0704-047 | HQ662213.1 | A | BD | - | --- | - | + | + | **6D** |
| 07-107 | HQ662212.1 | A | BD | - | --- | - | + | + | **6D** |
| 07-077 | HQ662211.1 | A | BD | - | --- | - | + | + | **6D** |
| 07-056 | HQ662210.1 | A | BD | - | --- | - | + | + | **6D** |
| Tw02-238 | HQ662209.1 | A | BD | - | --- | - | + | + | **6D** |

<sup>a</sup> wciP374F - wciP-r; <sup>b</sup>Del6Cwzy\_Fv2 - Del6Cwzy\_R; <sup>c</sup>wciN\_6AB\_F - wciN\_6AB\_R; <sup>d</sup>wciNbetaS1 – wciNbetaA2; <sup>e</sup>wciNbetaA1 – wciNbetaS2

**Figure 2.**
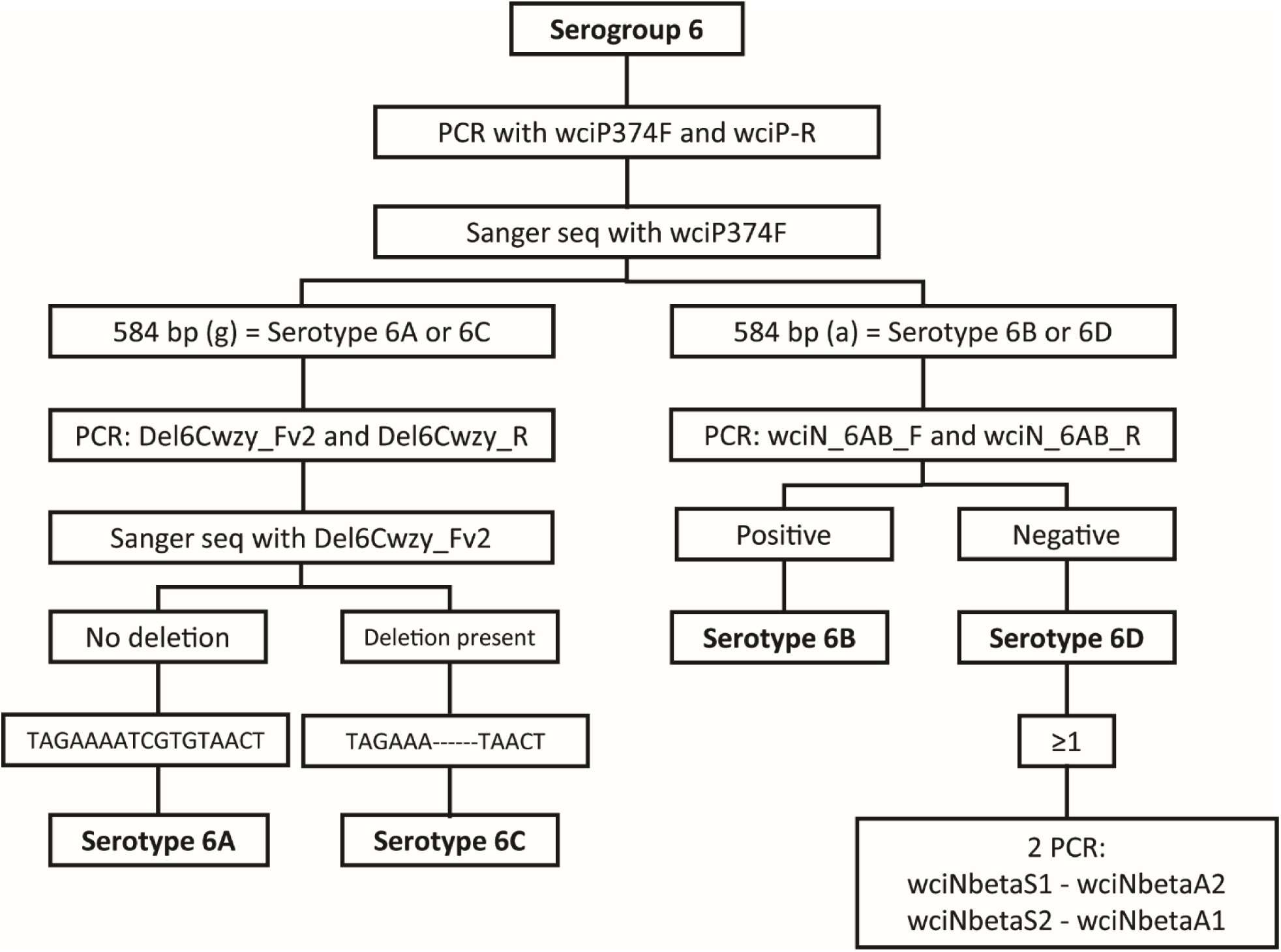
Schematic representation of serogroup 6 differentiation.

The sequence analyses performed for the target regions of the genomes showed that the regions where the primers anneal are highly conserved; thus, PCR-amplification is expected to be specific and reliable. The proposed protocol was tested in all CCUG strains identified as serogroup 6 by Quellung reaction and in the clinical isolates identified as serogroup 6. Similar results were obtained in 9 strains when the proposed protocol was tested, compared to the results obtained by Quellung reaction, except for CCUG 3114, which was previously described as 6A and reclassified as serogroup 6C.

## DISCUSSION

Correct identifications of *S. pneumoniae* strains are crucial for choosing the proper treatment options and for assessment of the burden of disease. As a general standard, routine culture-based identification of *S. pneumoniae* consists of bile solubility and optochin susceptibility tests (Richter et al., 2008). There is a percentage of isolates that give inconsistent results with optochin susceptibility and bile solubility and are referred to as ‘atypical’ pneumococci. In addition, similar biochemical properties are also present in a significant proportion of other closely related species of the Mitis group, such as *S. mitis*, and *S. pseudopneumoniae,* especially when samples are from respiratory sites (Keith et al., 2006; Rolo et al., 2013). The *in silico* analysis performed in this study showed that identification of *S. pneumoniae* by analysis of the *groEL* partial sequence was possible and reliable, whereas *S. pseudopneumoniae* and *S. mitis* could be misclassified as *S. pneumoniae*, suggesting that *groEL* is an unreliable marker for differentiating *S. pneumoniae* from its closest related species. In the studies of Glazunova *et al*. (2009) and Teng *et al*. (2002) (Glazunova et al., 2009; Teng et al., 2002), where *groEL* was proposed to differentiate *S. pneumoniae* from other species of the Mitis group, few strains of each species were used for the analysis; in our study, we included all the genomes sequences available in the database at the time the study was performed. These results point to the risk that partial gene sequence analysis may lead to misclassification, for example, due to horizontal transfer. Horizontal gene transfer and homologous recombination, involving *groEL*, between species most likely occurs, as has been suggested previously for *sodA* and *rpoB* genes (Varghese et al., 2017).

High degrees of horizontal gene transfer and homologous recombination (Chi et al., 2007; Jensen et al., 2016; Kilian et al., 2008) between *S. pneumoniae* and commensal viridans group streptococci have given rise to genotypic ambiguities between *S. pneumoniae* and closely related species, such as *S. mitis*, *S. pseudopneumoniae* and *S. oralis* (Kilian et al., 2008; Kilian et al., 2014; Whatmore et al., 2000). Multi-Locus Sequence Analysis (MLSA) for the Viridans group streptococci developed by Bishop *et al*. (2009) (Bishop et al., 2009) and core genome phylogenetic analyses (Jensen et al., 2016) are genome-based techniques that can differentiate the Viridans group streptococci to the species level. ANIb similarity determination is gaining relevance as a robust measure of relatedness between strains, with potential in confirming phylogenetic and taxonomic relationships of bacterial identification (Konstantinidis and Tiedje, 2005). An ANIb similarity threshold above 95%, with respect to species reference type strains is proposed to provide species-level identifications of given genomes (Kim et al., 2014; Richter and Rossello-Mora, 2009). In our study, the genome sequences of *S. pneumoniae* and *S. pseudopneumoniae* fulfilled the suggested thresholds, whereas only four of the *S. mitis* genome sequences fulfilled this criterion. However, cluster analyses, derived from determined all vs. all genome sequences ANIb similarity values, allowed discrimination of the different species, by clustering with respect to species type strains. The *in silico* DNA-DNA hybridization calculated with the GGDC was also inconclusive for identification of *S. mitis* genome sequences.

The recently described “Xisco” gene, which is detected by a single PCR, seems to be a good marker for the correct identification of *S. pneumoniae* and differentiation from the closely related species *S. pseudopneumoniae* and *S. mitis* (Salvà-Serra et al., 2017). Both in the *in silico* and *in vitro* analyses, the “Xisco” gene was present in all *S. pneumoniae* strains and absent in all genomes and strains of the non-pneumococcus Mitis group species. Other targets have been proposed to be specific for pneumococci the last decade, such as pneumolysin (*ply*) (McAvin et al., 2001), autolysin (*lytA*) (Corless et al., 2001), pneumococcal surface antigen A (*psaA*) (Morrison et al., 2000), and penicillin binding protein (*pbp*) (O’Neill et al., 1999), among others. However, the “Xisco” gene seems to be more robust and distinguishes *S. pneumoniae* from the other species of the Mitis group more reliably. Since recombination in the Mitis group may occur, it is potentially unreliable to use a single gene biomarker for identification of *S. pneumoniae*.

Identification of *S. pneumoniae* serogroup/serotype is important for surveillance of strains in disease carriage and for strategies in vaccine development (O’ Brien et al., 2009). For serotyping *S. pneumoniae*, the Quellung reaction method is considered the ‘gold standard‘, although it can be performed only on viable isolates, needs expertise and is expensive. Recently, molecular techniques, such as genotypic typing methods targeting serotype-specific regions of the *cps* genes, including multiplex PCR (Brito et al., 2003; da Gloria Carvalho et al., 2010; Jourdain et al., 2011; Pai et al., 2006; Richter et al., 2013) and multiplex real-time Q-PCR (Pimenta et al., 2013), have been described. These methods allow the detection of multiple serotypes but are still relatively laborious, considering than more that nearly 100 different serotypes are known today. Most of these methods were designed to be able to identify the serotypes that have been included in vaccines or which are most common in given geographic areas. However, in surveillance studies, replacement of vaccine serotypes by non-vaccine serotypes has been reported in regions where pneumococcal conjugate vaccines are implemented (Hicks et al., 2007; Weinberger et al., 2011), raising the necessity for simplified methods that allow detection of as many serotypes as possible, as well as recognition of newly-evolved serotypes,.

The recently described sequetyping technique by Leung *et al*. (2012), has the advantage of being able to detect a broad range of serotypes in one analysis. However, in our hands it was difficult to obtain adequate amplicons and, as pointed out by Leung *et al*., the size of the amplicon (1,061 bp) is too large for the current Sanger sequencing protocols. Therefore, we added two internal primers to amplify two fragments and sequence, enabling to obtain the whole *cpsB* sequence with good quality. This strategy allows distinction between the serotypes 18B and 18C, but not differentiation within the serogroups 6 (6A, 6B, 6C, and 6D) and 7 (7F and 7A). An advantage of sequetyping is that it can be based on data from the publicly available GenBank database, although, the nature of handling of data deposited with this database implies potential risk for incorrect assignment of serotype designations, as well as incorrect taxonomic classifications of strains. We confirmed that *cpsB* sequetyping gave correct serotype results by performing an *in silico* sequetyping of 329 available *S. pneumoniae* genomes and also a BLAST similarity analysis to further document the accuracy of the sequetyping method.

The utility of DNA-based methods for serotyping can be limited, due to inherent difficulties of differentiations within serogroups, which is of importance because available vaccines may include some, but not all, serotypes of a serogroup. Recently, PCR-based protocols for improved discrimination of serogroup 18 and serotypes 22F and 33F were described (Gillis et al., 2017; Tanmoy et al., 2016). Here we present a modified protocol for discriminating serogroup 6 serotypes, based on sequence analysis., The distinction is important, given the significant increase of pneumococcal infections of serotype 6C after introduction of conjugate vaccines.

The sequetyping was applied on 50 pneumococcal clinical isolates, which were also analysed by real-time Q-PCR and serotyping by an antiserum panel at the Public Health Agency of Sweden. The comparisons showed good agreement between the assays, similar to what was observed by Dube *et al*. (2015) (Dube et al., 2015). The results confirmed that sequetyping was able to detect also several non-vaccine serotypes. These genotypes were not detected by real-time Q-PCR because this assay identifies only those serotypes that are specifically targeted. However, the use of the sequetyping method is limited to single isolates due to difficulties to differentiate different serotypes when analysing the sequence chromatograms. In contrast, the real-time Q-PCR can be used with total DNA extracts from samples and is therefore, able to recognize the presence of multiple serotypes in a given sample.

In conclusion, the presence of the “Xisco” gene, genome sequence ANIb, *in silico* DNA-DNA hybridization and targeted *groEL* comparative sequence analyses are reliable methods for identification of pneumococci. Serotyping by using PCR-and DNA sequence-based methods is highly useful in cases where access to traditional methods is limited and when cultivation of isolates is negative. However, since *S. pneumoniae* and the related species of the Mitis group of Streptococcus undergo constant recombination, the use of the different techniques needs to be applied in order to verify the reliability of analyses.

## ACKNOWLEDGEMENTS

This work was supported by the European Commission: TAILORED-Treatment (project number 602860; www.tailored-treatment.eu). The Culture Collection University of Gothenburg (CCUG) is supported by the Department of Clinical Microbiology, Sahlgrenska University Hospital and the Sahlgrenska Academy of the University of Gothenburg. FS-S was supported by stipends for Basic and Advanced Research from the CCUG, through the Institute of Biomedicine, Sahlgrenska Academy, University of Gothenburg.

